# Complete mitochondrial genome of Glomeridesmus spelaeus (Diplopoda), a troglobitic species from Carajás iron-ore caves (Pará, Brazil)

**DOI:** 10.1101/228882

**Authors:** Gisele Lopes Nunes, Renato Renison Moreira Oliveira, Eder Soares Pires, Santelmo Vasconcelos, Thadeu Pietrobon, Xavier Prous, Guilherme Oliveira

**Affiliations:** Instituto Tecnológico Vale, Belém, PA, Brazil; Vale, Belo Horizonte, MG, Brazil

**Keywords:** Mitogenome, troblobite, iron-ore caves, *Glomeridesmus*

## Abstract

We report the complete mitochondrial genome sequence of *Glomeridesmus spelaeus,* the first sequenced genome of the order Gomeridesmida. The genome is 14,825 pb in length and encodes 37 mitochondrial (13 PCGs, 2 rRNA genes, 22 tRNA) genes and contains a typical AT-rich region. The base composition of the genome was A (40.1%), T (36.4%), C (15.8%), and G (7.6%), with an AT content of 76.5%. Our results indicated that *Glomeridesmus spelaeus* only distantly related to the other Diplopoda species with available mitochondrial genomes in the public databases. The publication of the mitogenome of *G. spelaeus* will contribute to the identification of troglobitic invertebrates, a very significant advance for the conservation of the troglofauna.

---

There are a large number of caves in Brazil, however only 7% are registered of a total estimated of 100,000 caves (Sallun Filho and Karmann, 2012; Auler and Piló, 2015). These registers are temporally validate, based in environmental studies conducted for mining operations, by National Center for Research and Conservation of Caves (CECAV - http://www.icmbio.gov.br/cecav). The Brazilian legislation considers caves as special environmental entities subjected to specific protective measures that are based on levels of relevance from Maximum to Minimum (Auler and Piló, 2015). Relevance level depends both on physical and biological attributes, among others, although the last are the most important in the evaluation of the cave relevance, and thus, studies on the cave biotas are of central importance (Jaffé et al., 2016). Nevertheless, there is a large knowledge gap on the identification of cave inhabitants, especially troglobites (those inhabiting exclusively the cave environment). As a consequence, stakeholders lack solid support for decisionmaking linked to the use of natural resources.

*Glomeridesmus spelaeus* is a troglobite species of the class Diplopoda found in iron ore caves in the Carajás region in the Amazon basin (Iniesta et al., 2012). *Glomeridesmus* are chilognath millipedes belonging to Glomeridesmida, a small order composed by 31 species (26 Glomeridesmidae and five Termitodesmidae) (Jeekel, 2003), which may be basal to pill millipedes and other millipedes (Hoffman et al., 1982). There is very limited knowledge on the troglobitic Glomeridesmida, with only two records described so far: *G. sbordonii* (Shear, 1974) from Grutas de Cocona at Teapa, Tabasco, Mexico and *G. spelaeus* (Iniesta et al., 2012) from Carajás iron-ore caves at Curionópolis, Pará, Brazil. Considering the scarcity of knowledge about Glomeridesmida species, we aim to provide genetic informations to troglobitic *G. spelaeus*, in order to contribute to a more comprehensive understanding of one of the least known arthropod orders.

A specimen was collected in a Carajás iron-ore cave (6S 04’ 52”, 50W 08’ 03”), Pará, Brazil (Figure1). Total genomic DNA was extracted using DNeasy Blood & Tissue Kit (Qiagen, Valencia, CA), following the recommended protocol for insect species. Paired-end libraries (2 × 75 bp) were built using QXT Sure Select (Agillent Technologies, San Diego, USA) and sequenced on the NextSeq platform (Illumina, San Diego, CA) generating a total of 29,527,776 high quality paired-ends. NOVOPlasty 2.6.3 (Dierckxsens et al., 2017) and Geneious R10 (Biomatters, Auckland, New Zealand) were used for a *de novo* assembly of the mitogenome, resulting in a single and circular mitochondrial genome. Annotations were carried out with the MITOchondrial genome annotation Server (MITOS) (Bernt et al., 2013).

**Figure 1.**
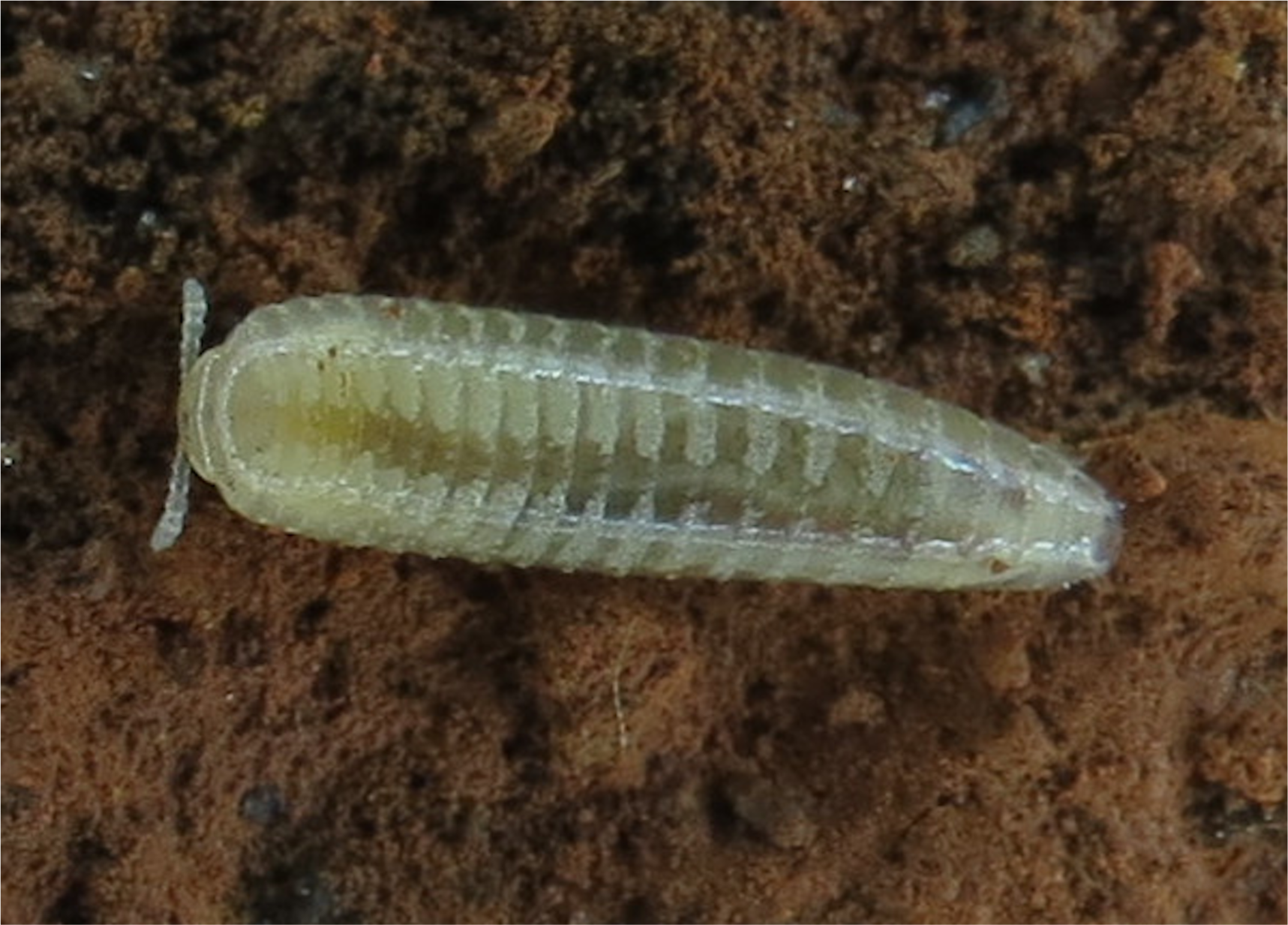
*Glomeridesmus spelaeus*.

The complete mitochondrial genome of *G. spelaeus* was 14,825 bp in size including 13 protein-coding genes (PCGs), two ribosomal RNAs and 22 transfer RNAs (Genbank accession MG372113) (Figure 2 and Table1). The overall nucleotide composition of the mitogenome was 40.1% of A, 15.8% of C, 7.6% of G, and 36.4% of T. The total AT and GC content was 76.5% and 23.5%, respectively. Most of the mitogenome genes are encoded on the H-strand, except for the regions nad1, nad4, nad4l nad5, trnC(gca), trnF(gaa), trnH(gtg), trnI(gat), trnL1(tag), trnL2(taa), trnP(tgg), trnQ(ttg), trnV(tac) and trnY(gta), which were encoded on the L-strand. The genes that were not initiated with ATG codon, started with ATA (cox1, nad4l, cob) and ATT (nad1, nad2, nad3, nad5, nad6 and atp8). All protein-coding sequences terminated with a conventional stop codon (TAA).

**Table 1.** Mitogenomes comparison of *Glomeridesmus spelaeus* and Sphaerotheriidae sp.

| Gene | Gene strand |  | Codons (start/stop) |  |
| --- | --- | --- | --- | --- |
|  | G.<br>spelaeus | Sphaerotheriidae sp. | G.<br>spelaeus | Sphaerotheriidae sp. |
| atp6 | + | + | ATG/TAA | ATG/TAA |
| atp8 | + | + | ATT/TAA | ATA/TAA |
| cob | + | + | ATA/TAA | ATG/TAG |
| cox1 | + | + | ATA/TAA | ATG/TAA |
| cox2 | + | + | ATG/TAA | ATG/TAA |
| cox3 | + | + | ATG/TAA | ATG/TAA |
| nad1 | - | - | ATT/TAA | ATG/TAA |
| nad2 | + | + | ATT/TAA | ATT/TAA |
| nad3 | + | + | ATT/TAA | ATA/TAG |
| nad4 | - | - | ATG/TAA | ATG/TAA |
| nad4l | - | - | ATA/TAA | ATG/TAA |
| nad5 | - | - | ATT/TAA | ATT/TAA |
| nad6 | + | + | ATT/TAA | ATT/TAA |
| rrnL | - | - |  |  |
| rrnS | - | - |  |  |
| trnA(tgc) | + | + |  |  |
| trnC(gca) | - | - |  |  |
| trnD(gtc) | + | + |  |  |
| trnE(ttc) | + | + |  |  |
| trnF(gaa) | - | - |  |  |
| **trnG(tcc)** | + | 0 |  |  |
| trnH(gtg) | - | - |  |  |
| trnI(gat) | - | + |  |  |
| **trnK(ctt)** | 0 | + |  |  |
| trnL1(tag) | - | - |  |  |
| trnL2(taa) | - | - |  |  |
| trnM(cat) | + | + |  |  |
| trnN(gtt) | + | + |  |  |
| trnP(tgg) | - | - |  |  |
| trnQ(ttg) | - | - |  |  |
| trnR(tcg) | + | + |  |  |
| trnS1(gct) | + | + |  |  |
| trnS2(tga) | + | + |  |  |
| trnT(tgt) | + | + |  |  |
| trnV(tac) | - | - |  |  |
| trnW(tca) | + | + |  |  |
| trnY(gta) | - | - |  |  |

**Figure 2.**
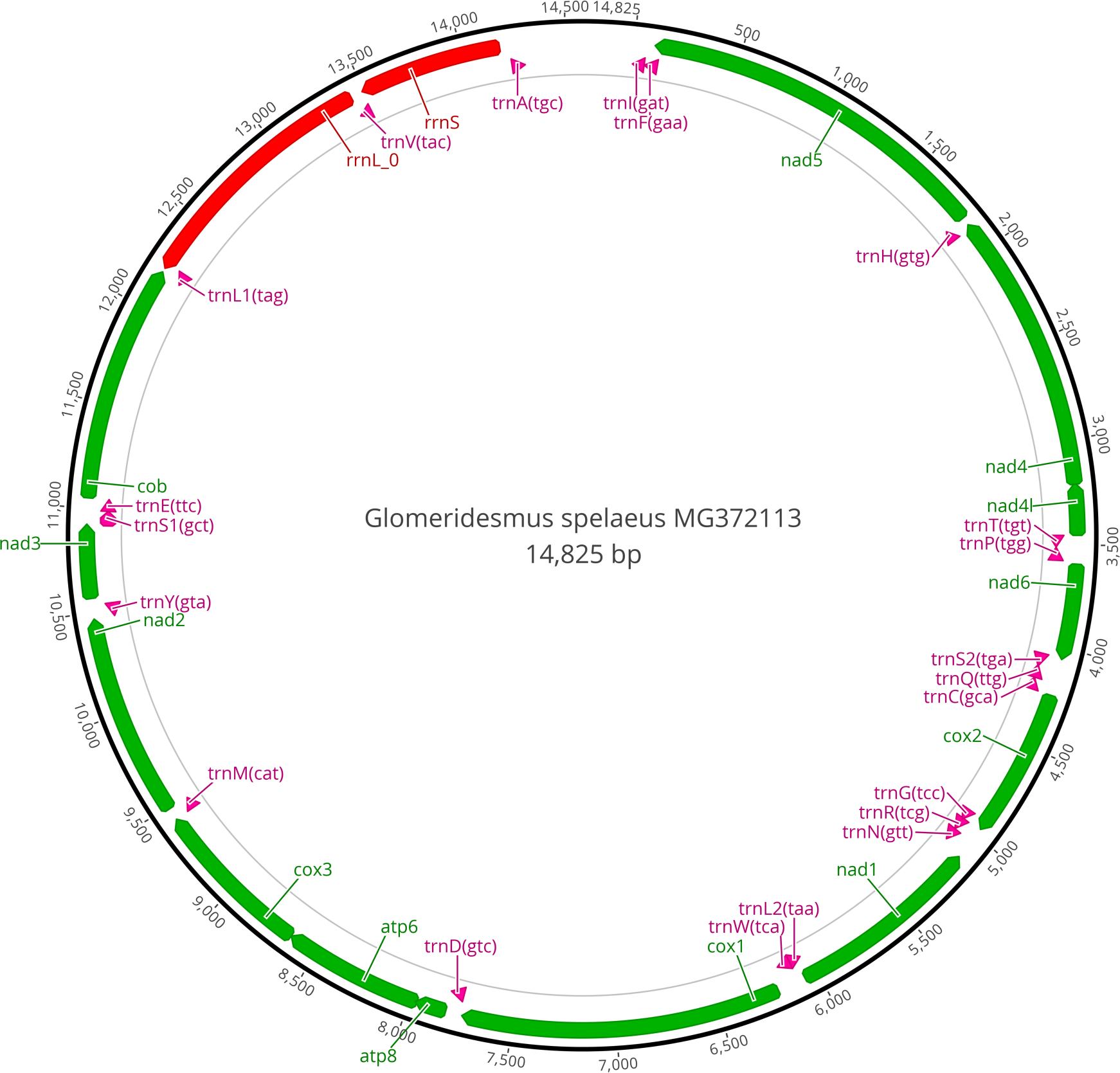
Complete mitochondrial genome of *Glomeridesmus spelaeus.* The colors green, pink and red correspond to protein coding gene, tRNA gene, rRNA gene, respectively.

In addition, we performed a phylogenetic analysis based on all mtDNA genes (except for those either missing or duplicated in at least one genome) from all available mitochondrial genomes of the Diplopoda species that were available in Genbank *(Abacion magnum, Anaulaciulus koreanus, Antrokoreana gracilipes, Appalachioria falcifera, Asiomorpha coarctata, Brachycybe lecontii*, Sphaerotheriidae sp. and *Xystodesmus* sp.) and two Chilopoda mtDNA genomes as outgroup *(Cermatobius longicornis* and *Scutigera coleoptrata*) (Figure 3). All used genes were aligned separately in MAFFT 7.3 (Katoh and Standley, 2013), using the algorithm *Auto*, and then concatenated in a single matrix. Afterwards, the phylogenetic tree was obtained through the maximum likelihood approach implemented in RAxML 8.2 (Stamatakis, 2014) using rapid bootstraping with 1,000 replicates. The phylogenetic results supported that *G. spelaeus* was more closely related to an unknown species belonging to the family Sphaerotheriidae, although only distantly related. A genetic comparison using the complete mitogenomes of *G. spelaeus* and Sphaerotheriidae sp. is represented in Table 1. All genes are present in both mitogenomes except trnK (present only in Sphaerotheriidae sp.) *and* trnG (present only in *G. spelaeus).* We observed no difference on gene transcription strand between the two species, except for the trnI gene with L strand in *G. spelaeus* and H strand in Sphaerotheriidae. Stop and start codons were different when compared the same gene groups, except for atp6, cox2, cox3, nad2, nad4, nad5, nad6 that show the same genetic code for both initiation and termination of the translation process (Table 1).

**Figure 3.**
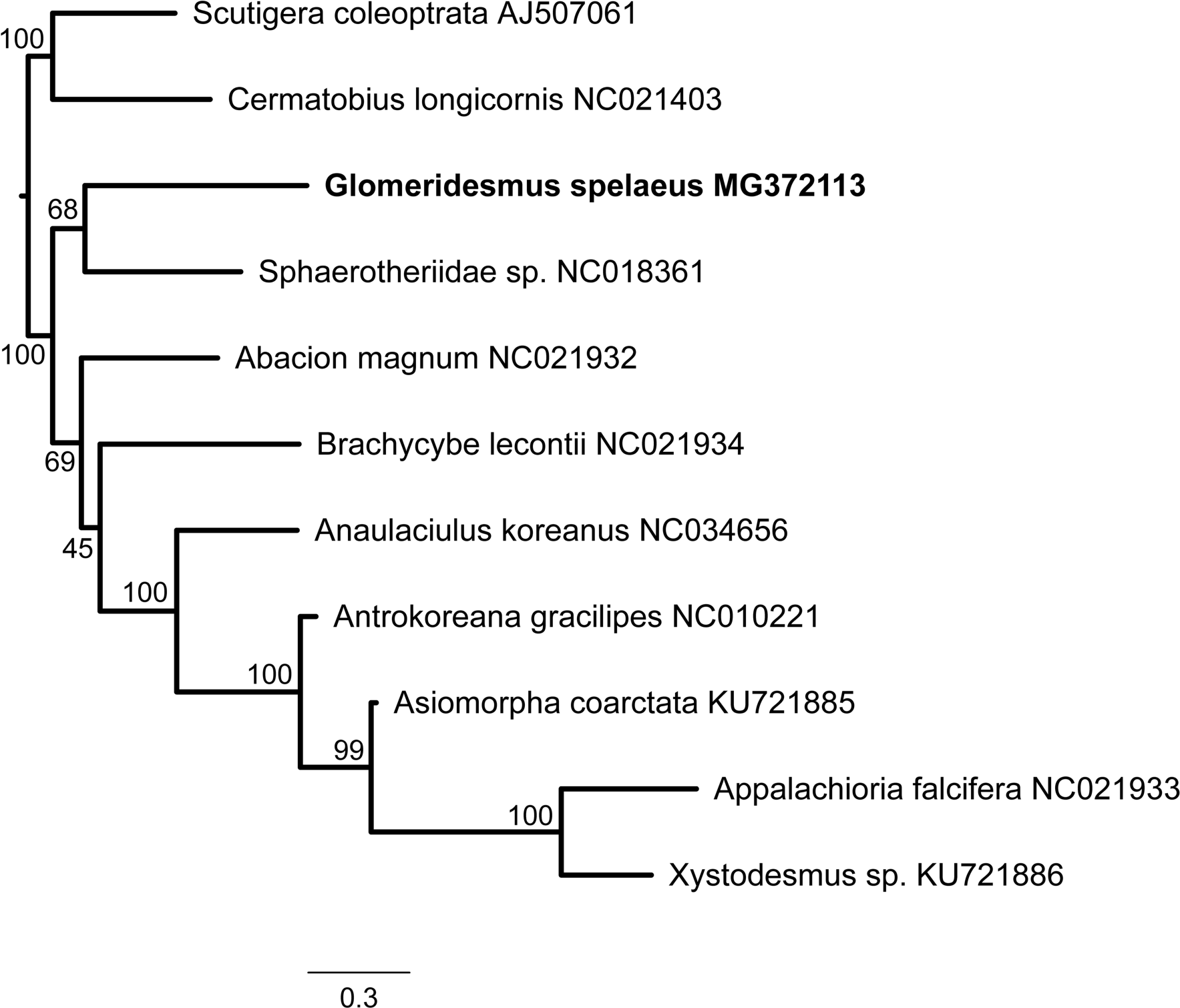
Phylogenetic tree of *Glomeridesmus spelaeus* and Diplopoda species. A Maximum likelihood tree was constructed using *GTR-Gamma* model with 1000 bootstrap replicates. Genbank accession number for each sequence are shown with the species name.

There is still a remarkable gap of information for the order Glomeridesmida in public databases. Our data provides the first genetic information to order Glomeridesmida. Research including troglobitic species are essential to understand both speciation and evolution process of these organisms that live exclusively in caves. Genetic data on such species will be central for the determination of the biodiversity present in the ferruginous caves of Carajás in addition to contribute to the unraveling of the evolutionary patterns of the genus *Glomeridesmus*.

## Acknowledgments

This work was funded by Vale. The funders had no role in the study design, data collection and interpretation, or the decision to submit the work for publication. This work, including the efforts of Renato Oliveira funded by MCTI | Conselho Nacional de Desenvolvimento Científico e Tecnológico (CNPq) (459913/2014-0), Gisele Lopes Nunes funded by CAPES (88887.141259/2017-00), Eder Soares Pires funded by CAPES (88887.141260/2017-00). Guilherme is a CNPq fellow (307479/2016). This work was funded by Vale (TrogloGen Project). The authors declare no conflict of interest.

